# Detection of live breast cancer cells in brightfield microscopy images containing white blood cells by image analysis and deep learning

**DOI:** 10.1101/2021.11.04.467215

**Authors:** Golnaz Moallem, Adity A. Pore, Anirudh Gangadhar, Hamed Sari-Sarraf, Siva A. Vanapalli

## Abstract

**Significance:** Circulating tumor cells (CTCs) are important biomarkers for cancer management. Isolated CTCs from blood are stained to detect and enumerate CTCs. However, the staining process is laborious and moreover makes CTCs unsuitable for drug testing and molecular characterization.

**Aim:** The goal is to develop and test deep learning (DL) approaches to detect unstained breast cancer cells in bright field microscopy images that contain white blood cells (WBCs).

**Approach:** We tested two convolutional neural network (CNN) approaches. The first approach allows investigation of the prominent features extracted by CNN to discriminate cancer cells from WBCs. The second approach is based on Faster Region-based Convolutional Neural Network (Faster R-CNN).

**Results:** Both approaches detected cancer cells with high sensitivity and specificity with the Faster R-CNN being more efficient and suitable for deployment. The distinctive feature used by the CNN used to discriminate is cell size, however in the absence of size difference, the CNN was found to be capable of learning other features. The Faster R-CNN was found to be robust with respect to intensity and contrast image transformations.

**Conclusions:** CNN-based deep learning approaches could be potentially applied to detect patient-derived CTCs from images of blood samples.

## 1. Introduction

Breast cancer is one of the leading causes of death with one in eight women expected to be diagnosed with invasive breast cancer in the United States [1]. An estimated 284,200 new cases are expected to be diagnosed in 2021 alone with about 44,130 deaths due to breast cancer [2]. A majority of deaths in breast cancer patients is due to metastasis, where CTCs shed from the primary tumor, spread to distant organs and form secondary tumors [3, 4]. Isolating, enumerating, and characterizing these CTCs are therefore valuable means to monitor disease progression and manage treatment therapies for cancer patients [5–9].

The CTCs are present in exceedingly low counts in peripheral blood (1-100 CTCs/*ml*) [10] , which has resulted in significant efforts to develop methods for isolating and characterizing these rare cells. Current CTC isolation techniques may be classified into two broad categories: affinity-based (labeled) and label-free techniques. Technologies based on the former approach rely on the interaction of cell-surface receptors on CTCs and specific antibodies, leading to the isolation of only the CTCs that bind to these antibodies [11–14]. In contrast, label-free approaches isolate CTCs based on physical characteristics, e.g., size [15–17], deformability [18], dielectric properties [19, 20] and density [21].

Despite the development of numerous separation methods for isolating CTCs from whole blood, the enriched CTC samples are almost always accompanied by a large number of WBCs [22–27]. The presence of these background nucleated cells makes it necessary to use molecular markers and immunostaining to distinguish CTCs from WBCs. However, the immunostaining process kills CTCs; thus, making it impossible to use them in downstream assays such as in vivo studies, expansion assays or single-cell transcriptomics, all of which require the CTCs to be alive and functional [28–30]. Thus, there is a need for staining-free approaches that can enumerate live CTCs in a background of WBCs.

Current approaches to detecting stained CTCs range from manual scoring to techniques based on image processing and machine learning. There is growing interest in using machine learning to detect and classify cells in patient blood samples as it eliminates drawbacks of manual scoring and threshold-based object detection [31–33]. For example, recently, Zeune *et al*. [34] developed a deep learning network to automatically identify and classify fluorescently stained CTCs obtained from the blood of metastatic cancer patients with a reported accuracy of over 96%. Deep learning networks are attractive as they do not require any feature engineering but are capable of automatically discovering the best set of features that discriminate the cells of interest and can directly identify them from raw input images [35].

Given the success of deep learning models to identify CTCs in stained images, new efforts are emerging to implement similar approaches in unstained samples. However, since CTCs are quite heterogeneous and are present in low counts in patient samples, generating ground-truth data is challenging. Thus, as a first step, deep learning (DL)-based frameworks are being pursued where in vitro cancer cells are mixed with blood cells and the efficacy of deep learning models are being investigated to detect live cancer cells in a background of blood cells [36–39] As shown in Table 1, limited studies have been conducted on DL-based frameworks for cancer cell identification from a background of blood cells, with images acquired using quantitative phase, dark field, and bright field microscopy. A notable study of this kind is the study by Wang *et al.*, who demonstrated detection of live CTCs in the blood of renal cancer patients using a deep convolutional neural network (CNN) and reported an accuracy of 88.6% [36] .

**Table 1:** Summary of studies that applied deep-learning methods for the identification of unstained *in vitro* cancer cells in a background of blood cells. The performance measures shown for our work correspond to the Faster R-CNN model.

| *Study* | *Cancer cells* | *Background cells* | *Test images* | *Ground truth images* | *Classification/ Detection Model used* | *Accuracy/ Sensitivity/ Specificity/ Precision* |
| --- | --- | --- | --- | --- | --- | --- |
| Chen *et al.* 2016 [37] | Colon (SW-480) | T-cells (WBC subset) | Quantitative phase images | Quantitative phase images | DNN classifier trained on AUC | 95.5 %<br>Not Reported<br>Not Reported<br>Not Reported |
| Ciurte *et al.* 2018 [38] | Breast (Hs 578T, MCF-7), Colon (DLD-1) | Blood cells (RBCs + WBCs) | Darkfield images | Fluorescent images | Adaboost classifier | Not Reported<br>92.87 %<br>99.98 %<br>Not Reported |
| Wang *et al.* 2020 [36] | Colon (HCT-116) | WBCs | Brightfield images | Fluorescent images | ResNet50 CNN classifier | 88.6% for patient blood sample images and 97% for cultured cell images<br>Not Reported<br>Not Reported<br>Not Reported |
| This work | Breast (MCF-7) | WBCs | Brightfield images | Fluorescent images | Custom CNN classifier, Faster R-CNN detection model | 99.45%<br>99.1 %<br>Not applicable<br>99.8 % |

In this study, we utilize CNN approaches to demonstrate an automatic, sophisticated detection model for MCF-7 breast cancer cells in brightfield images of mixed population samples containing MCF-7 cells and WBCs. We develop two CNN models: (i) the first approach referred to as “decoupled cell detection” allows us to investigate the prominent features extracted by CNN to discriminate cancer cells from white blood cells. To the best of our knowledge, the features CNN uses to discriminate cells have not been explained by previous studies. (ii) the second approach is based on Faster Region-based Convolutional Neural Network [40] referred to as the Faster R-CNN, which is more efficient and therefore suitable for deployment. Additionally, we introduce a novel automatic technique to generate the training set for Faster R-CNN approach. This algorithm employs the brightfield image and the associated fluorescent images as ground truth. Additionally, we investigate the effect of image transformation on the classification accuracy to assess how well the deep learning model performs when variability is introduced in image quality. Finally, we discuss avenues for moving beyond the model system studied here and the challenges that need to be addressed to apply deep learning techniques to live CTC detection in patient blood samples.

## 2. Sample preparation, image acquisition and data sets

### Cell Culture

The breast cancer cell line MCF-7 was obtained from American Type Cell Collection (ATCC, Manassas, VA). MCF-7 cells were cultured in Dulbecco’s Modified Eagle Medium (DMEM, Gibco, Gaithersburg, MD) containing 10% Fetal Bovine Serum (FBS, Gibco, Gaithersburg, MD), 1% Penicillin/ Streptomycin (Gibco, Gaithersburg, MD) and 1% sodium pyruvate (Gibco, Gaithersburg, MD). The cell culture media was made according to the manufacturer’s protocol. The cells were incubated at 37°*C* in a 5% CO_2_ environment.

### MCF-7 Cell Staining

The cells were labelled with CellTracker™ Green CMFDA (Invitrogen™, Waltham, MA). The dye was prepared according to the manufacturer’s protocol to make a stock solution of 10 mM. A working solution of 1 μM was made by diluting the stock solution in serum free media. This working solution was added to the tissue culture flask containing cells followed by an incubation step for 45 minutes at 37°*C*. After incubation, the cells were washed with PBS three times to get rid of any excess dye. The cells were further trypsinized, resuspended in fresh media and stained with the nuclear stain DAPI (4′,6-diamidino-2-phenylindole, Molecular Probes, Eugene, OR). A working concentration of 100,000 cells/mL was used for the experiments.

### WBC Isolation and Staining

Normal Human Whole Blood was ordered from BioIVT (Hicksville, NY) for isolation of WBCs (White Blood Cells) from whole blood. 1 mL of whole blood was lysed using ACK lysing buffer (Quality Biological, Gaithersburg, MD) for isolation of WBCs. WBCs were stained with the nuclear stain DAPI. A working concentration of 3.6 × 10^6^ cells/mL was used for the experiments.

### Pure Cell Population Images

Slides of MCF-7 cells were prepared by adding 10 μL of working solution of MCF-7 cells on the cover glass (Richard-Allan Scientific, Kalamazoo, MI) equipped with 10 mm × 24 mm spacers on the edges. This solution was then sandwiched using another cover glass and imaged using Olympus IX81 microscope (Massachusetts, USA). The microscope is equipped with a Thorlabs automated stage (New Jersey, USA) and a Hamamatsu digital camera (ImagEM X2 EM-CCD, New Jersey, USA) controlled by SlideBook 6.1 (3i Intelligent Imaging Innovations Inc., Denver, USA). Brightfield and fluorescence images were acquired using 20x objective (0.8 μm/pixel, 512 × 512 pixels). Exposure times between 30-200 ms were used for image acquisition.

Similarly, slides of WBCs were prepared by adding 10 μL of working solution of WBCs on the sandwiched cover glass equipped with spacers. Brightfield and fluorescence images were acquired using 20x objective (0.8 μm/pixel, 512 × 512 pixels) and the fluorescent filters DAPI and FITC. Exposure times use for the WBC pure cell training set image acquisition was 30-100 ms.

### Mixed Cell Population Images

Slides for imaging of mixed cell populations were prepared by mixing 5 μL of WBC working solution with 5 μL of MCF-7 working solution. This 10 μL solution was added to the cover glass and imaged using the similar sandwich method as described previously [41, 42] . Brightfield and fluorescent (DAPI, FITC) image acquisition was done at 20x magnification. Imaged were acquired between 30-200 ms exposure times. Cell body and nucleus of MCF-7 cells were fluorescently labelled using CellTracker™ Green CMFDA and DAPI respectively, while those of WBCs were only labelled with DAPI, enabling distinction of the two cell types under fluorescence imaging.

### Image data sets

The brightfield images of MCF-7s and WBCs were accompanied by ground-truth DAPI and FITC fluorescent images for exploration, training and evaluation of deep-learning models used in this study. We acquired 224 pure MCF-7 cell images and 21 pure WBC images totaling 1131 MCF-7 cells and 11518 WBCs. For the mixed cell population, we acquired 368 images which contained 2769 MCF-7 cells. Fig. 1(A, B) shows examples of pure cell and mixed cell brightfield images along with their associated fluorescent counterparts.

**Figure 1.**
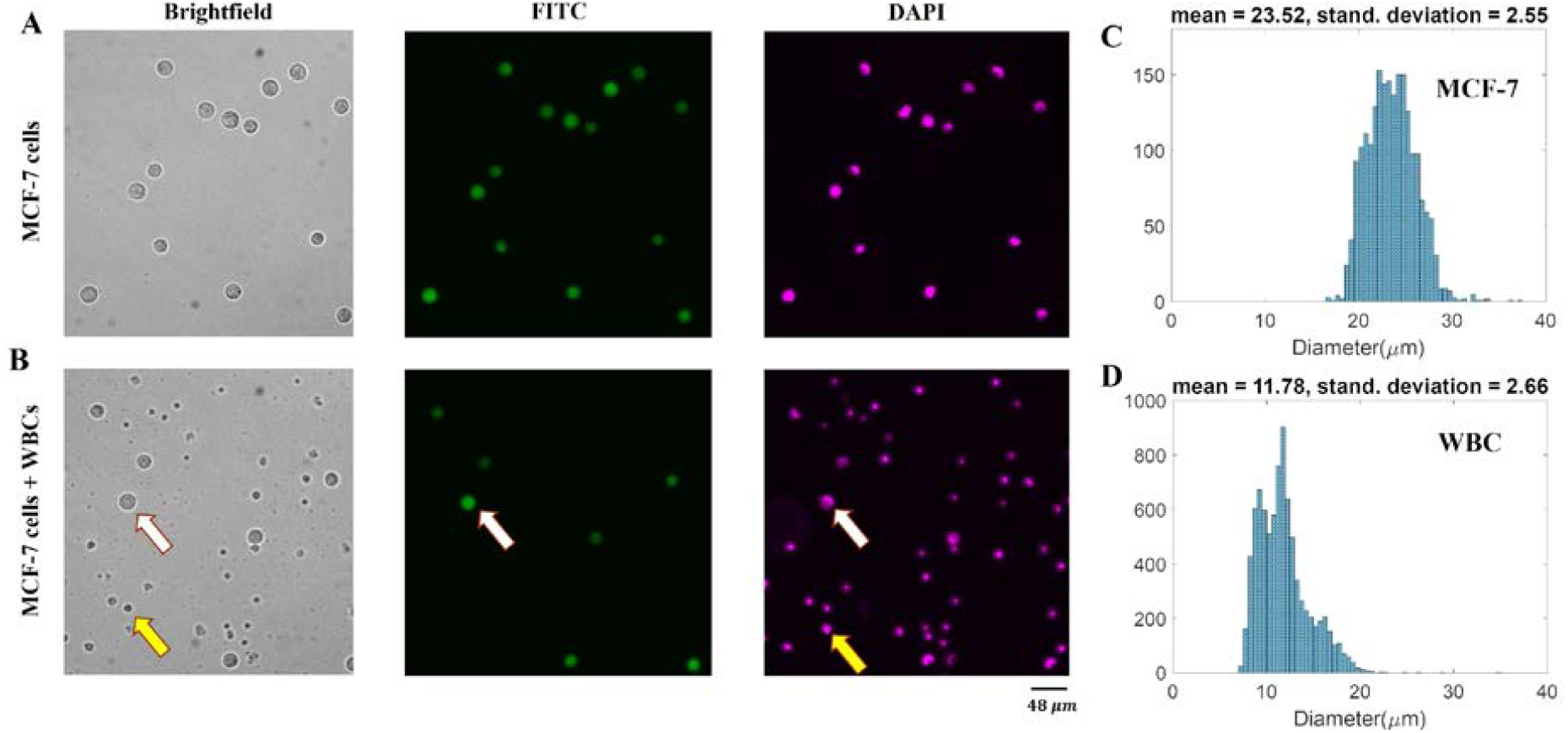
Representative images and size distribution of MCF-7 cancer cells and white blood cells (WBCs). (**A**) Brightfield and the corresponding fluorescent images of live MCF-7 cells. (**B**) Brightfield and corresponding fluorescent images of a mixed-cell population of MCF-7s and WBCs. White arrows denote an MCF-7 cell which can be co-localized in the ground-truth fluorescent FITC and DAPI images. Yellow arrows denote a WBC which can be co-localized in the DAPI image but not in the FITC image. (**C**) Size distribution of MCF-7 cells, n =2071. (**D**) Size distribution of WBCs, n = 8575.

## 3. Results and Discussion

In this study, we investigated two CNN-based approaches to detect MCF-7 cancer cells from a background of WBCs. In the first approach referred to as decoupled cell detection, all cells are localized using the Maximally Stable Extremal Regions (MSER) algorithm [43], and then classified using a trained CNN, *i.e.* the cell localization and identification tasks are carried out in two distinct steps. The benefit of the decoupled approach is that it allows investigation of the distinctive features that convolutional layers extract to distinguish MCF-7 cells and WBCs.

In the second approach, we detect MCF-7 cells using a Faster R-CNN model. In this approach, the cell localization and identification tasks are integrated without having separate access to the output of the localization step or the input of the classification step. As a result, it is difficult with Faster R-CNN model to understand the features being used to discriminate cells. However, due to its speed and efficiency of execution, the Faster R-CNN approach is better suited for deployment.

Below we discuss the results from our investigations with these two approaches to detect MCF-7 cells in a background of WBCs. We note that the network performance is reported in terms of accuracy, sensitivity, and precision [44]. Here, true positives (TPs) were defined as the number of detected MCF-7 cells. False negatives (FNs) were defined as the number of MCF-7 cells missed in the detection procedure, and false positives (FPs) were defined as the number of detections that did not correspond to MCF-7 cells.

### A. Decoupled Cell Detection

In this approach, the MCF-7 cells are detected in two decoupled modules: cell localization and cell identification modules. To detect the MCF-7 cells in a given brightfield image: (1) the brightfield image is fed to the localization module to localize the cells using the MSER algorithm, and (2) tiles of localized single cells are provided to the identification module to distinguish MCF-7 cells employing a trained CNN. As noted earlier, one of the purposes of developing this framework was to study the distinctive features that convolutional layers extract to distinguish MCF-7 cells and WBCs. Therefore, it was essential to have access to the input of the CNN classifier in order to solely study and evaluate its performance.

To develop the decoupled cell detection approach, we first designed and tested a shallow CNN using tiles of individual MCF-7 cells and WBCs. Subsequently, we evaluated the prominent cell features that allows successful classification with this trained CNN. Next, we optimized the CNN architecture to improve the detection of MCF-7 cells. Finally, we applied the decoupled cell detection approach to mixed cell images, *i.e.*, individual images that contained a mixture of MCF-7 cells and WBCs. The results from this systematic investigation are discussed below.

#### A shallow CNN efficiently discriminates MCF-7 cells from WBCs

To generate the training and testing set for the shallow CNN, the MSER algorithm was applied to localize the cells in the ‘pure’ brightfield images, *i.e.*, images containing only MCF-7 cells or WBCs. For each localized cell, a 36 × 36 tile was cropped with the cell at the center. The tile size was determined based on the MCF-7 estimated cell size distribution (Fig. 1C) and image resolution. This process ensured that both WBCs and MCF-7 cells were contained entirely in the cropped tiles. The tiles were carefully inspected, verified, and labeled manually employing the FITC and DAPI masks. Those tiles with a complete WBC located in the center of the corresponding tile were labeled ‘WBC’. Similarly, tiles with a complete MCF-7 cell positioned in the central region of the corresponding tile were tagged ‘MCF-7’. The training set was formed by creating balanced classes of WBC and MCF-7 tiles, each class comprising 1190 tiles. The testing set contained 510 tiles per class.

We initially designed and trained a shallow CNN with two convolutional layers followed by a fully connected layer to classify the tiles into ‘WBC’ and ‘MCF7’ categories. The convolutional layers had 5 × 5 kernel sizes followed by a 2 × 2 pooling layer to minimize the encoding depth due to the relatively small size of the tiles. The trained CNN was found to achieve satisfactory performance with a training accuracy of 99.51% and a test accuracy of 99.54%. These performance metrics justified the use of the relatively shallow architecture of the CNN as it achieved a high accuracy level without overfitting. In addition, the class probability histograms showed the network’s high confidence level of above 99% on classification decisions. We concluded that the trained CNN successfully performed the discrimination task.

#### Identifying the discriminatory cellular features

We investigated the cellular features that enabled the successful classification by the shallow CNN. Since in Fig. 1(C, D) we observed that the mean diameter of MCF-7 cells is approximately twice as large as the mean diameter of WBCs, we hypothesized that cell size could be a distinguishing feature. To test this hypothesis, we made the size of WBCs similar to MCF-7 cells. This was achieved by doubling the size of tiles, which doubled the size of the WBCs contained in them, and then they were cropped to make them compatible with the network input. When we tested the trained CNN on the newly generated WBC tiles, we observed a deterioration in the classification accuracy to 61%, which confirmed that the trained CNN indeed relied on the size feature to make the classification decision.

Next, we hypothesized that in addition to cell size, the shallow CNN could extract other geometric and photometric features such as texture of the cell to perform the classification task successfully. To test this hypothesis, we eliminated the size difference between the cell types and trained our CNN in two ways. (i) We doubled the size of training ‘WBC’ tiles and used them together with the ‘MCF7’ tiles to train the CNN. In doing so, we achieved a 99.33% training accuracy and a 98.14% testing accuracy. (ii) Alternatively, training ‘MCF7’ tiles were halved in size and combined with the ‘WBC’ tiles to train the CNN. This resulted in a 98.32% training and a 96.86% test accuracy. The high training and testing accuracy suggest that although size is a prominent discriminatory feature used by the designed CNN, the network does utilize other features such as texture to perform cell classification with a high, albeit reduced, level of accuracy if necessary. This result helps explain the model’s ability to distinguish the two cell types, even when there is an overlap in the cell size distribution (Fig. 1C).

#### Optimizing the CNN architecture

The training was carried out on two different CNN architectures, one with two and the other with three convolutional layers while keeping the kernel sizes as 5 × 5 and pooling layer as 2 × 2. To boost the true positive detection rate of the CNN, we tuned the MSER parameters to maximize the localization of MCF-7 cells at the expense of a proportional increase in the overall number of tiles that were output by the MSER module. The classification task was then recast as a two-class problem in which one class represented MCF-7 cells, and the second represented all non-MCF-7 objects that included WBCs and debris. The results showed that the CNN with three convolutional layers had testing sensitivity and precision of 98.8%, outperformed the CNN with two layers. We therefore used the CNN with three convolution layers in subsequent studies for classification in the decoupled cell detection approach.

#### Validation of the decoupled cell detection approach on mixed-cell images

So far, our investigations were focused on pure cell images to develop and optimize a shallow CNN and identify cellular features that enable classification. Next, we validated the decoupled cell detection approach on images containing both MC7s and WBCs, as shown in Fig. 2.

**Figure 2.**
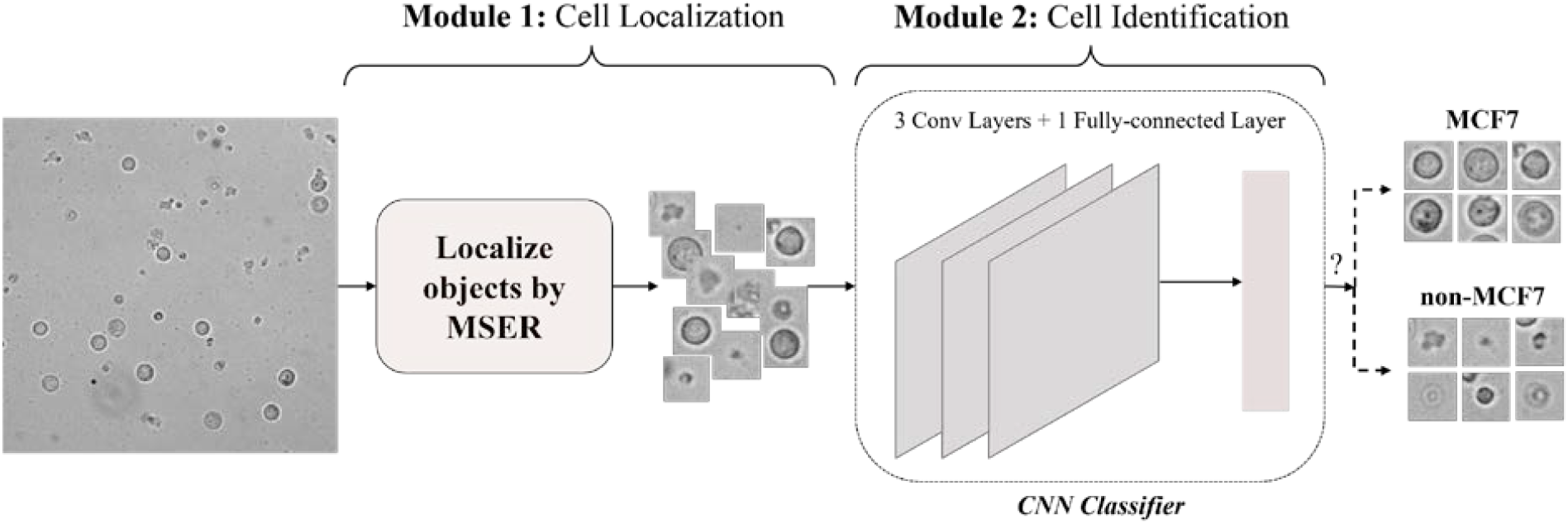
The framework of the decoupled cell detection approach. As shown, the cell detection is performed in two modules: cell localization and cell identification. The localization module localizes the cells present in the input brightfield image. The CNN-based identification module then identifies the MCF-7 cells among the localized cells.

In the localization module, the MSER algorithm is applied to the input image to localize the objects. One of the parameters of the MSER algorithm is the size range of the regions it detects. This parameter was set based on the lower and upper bounds of the MCF-7 size distribution to maximize the localization of MCF-7 cells. Specifically, the size range was set to [lower bound-5%, upper bound+5%]. Outputs from the localization module are 36 × 36 tiles centered on the corresponding objects. The tile size was calculated based on the upper bound of the MCF-7 cell size distribution and image resolution.

In the identification module, the cropped tiles are fed to the trained CNN model that, as described above, consists of three convolutional layers and a fully connected layer. The convolutional layers have a 5 × 5 kernel size, followed by a 2 × 2 pooling layer. The fully connected layer outputs a 2-dimensional vector containing the ‘MCF7’ and ‘non-MCF7’ class prediction scores.

The training and testing set generation workflow for developing the decoupled cell detection framework is illustrated in Fig. 3. In this workflow, first, MSER was applied to each input image, and tiles of localized objects were generated. Afterward, the cropped tiles were manually categorized based on their corresponding signatures in DAPI and FITC masks. Every localization depicting a signature in both DAPI and FITC masks was labeled as ‘MCF7’ since cancer cells were live stained with both a nucleus (DAPI) and a cell body (FITC) marker, while tiles lacking any signature in the FITC mask were annotated as ‘non-MCF7’.

**Figure 3.**
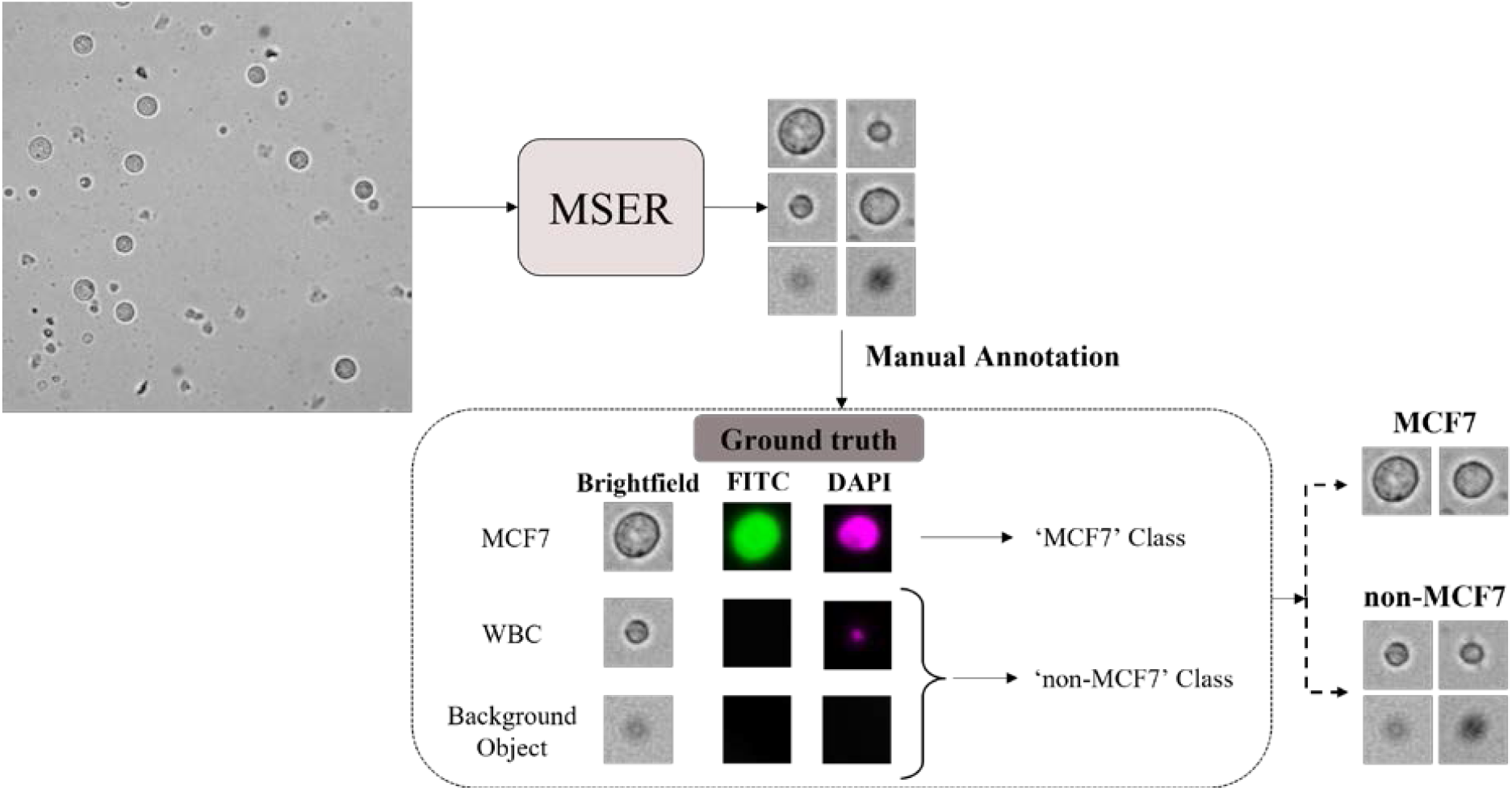
Training and testing set generation strategy for developing the framework of the decoupled cell detection approach. As demonstrated, the objects in the mixed population image are localized employing the MSER algorithm. Then, the generated tiles of the localized objects are manually categorized based on their corresponding signatures in the FITC and DAPI filters.

We used the described training dataset to generate the training tiles, resulting in 2,760 ‘MCF7’ tiles and 2,760 tiles in the ‘non-MCF7’ class. The CNN was trained using the ADAM optimizer for 35 epochs. The training process took 74 seconds on an Nvidia TITAN RTX. The trained classification CNN achieved 98.99% training sensitivity and 99.8% training precision.

#### Performance of the decoupled cell detection approach on mixed-cell images

We evaluated the decoupled approach on the testing set using three performance metrics: sensitivity, precision, and average execution time per image as shown in Table 2. Overall, the trained CNN performed with 95.5 % accuracy, 95.3% sensitivity and 99.8% precision, which indicates its ability to successfully achieve the MCF-7 identification task. Fig. 4 illustrates representative examples of TPs, FPs, and FNs observed during the analysis. In the decoupled approach, MCF-7 cells localized and correctly classified were counted towards TPs (Fig. 4A). The combination of (i) the MCF-7 cells not localized (Fig. 4B) and (ii) the MCF-7 tiles labeled as non-MCF7 (Fig. 4C) were counted as FNs. Lastly, the non-MCF7 tiles categorized into the MCF-7 class were considered as FPs (Fig. 4D).

**Table 2:** The performance statistics of the two CNN approaches used in this study.

| Approach | TPs | FNs |  | FPs |  | Sensitivity | Precision | Average Inference time per image |
| --- | --- | --- | --- | --- | --- | --- | --- | --- |
|  |  |  |  | Non-MCF7 Objects | Extra Detections |  |  |  |
| *Decoupled Cell Detection* | 1,869 | Module1 | Module 2 | 3 | 1 | 95.3% | 99.8% | 0.6 seconds |
|  |  | 71 | 22 |  |  |  |  |  |
| *Faster R-CNN Cell Detection* | 1,945 | 17 |  | 4 | 0 | 99.1% | 99.8% | 0.3 seconds |

**Figure 4.**
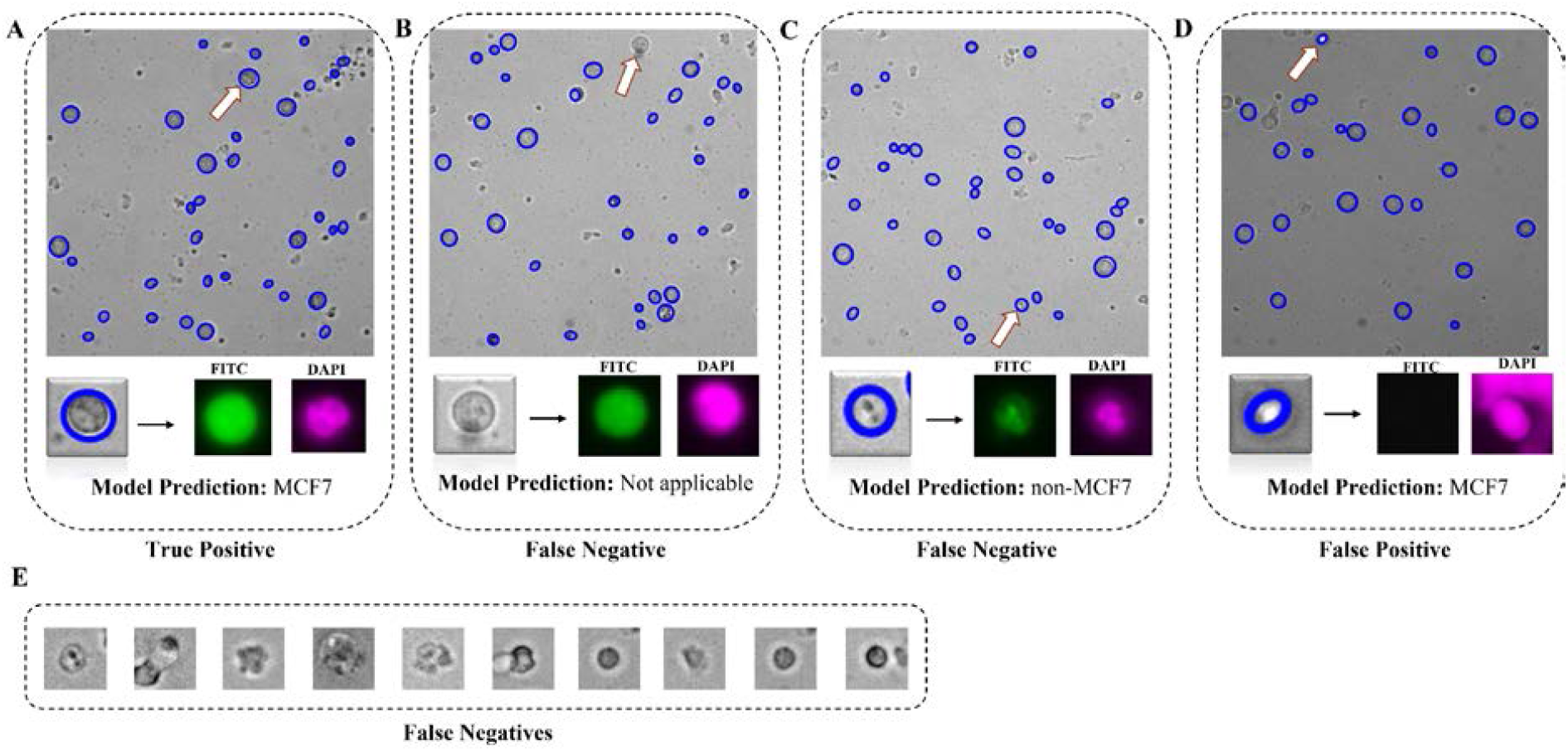
Representative examples of true positive (TP), false positive (FP) and false negative (FN) outcomes from the decoupled cell detection approach. (**A**) TPs are those MCF-7 cells that are localized and correctly identified as MCF-7s by the model. (**B**) FNs are MCF-7 cells that are not localized and therefore the model is not able to reach the identification stage (**C**) FNs can also be MCF-7 cells that are localized, but incorrectly classified by the model as non-MCF-7. (**D**) FPs are the non-MCF-7 objects that are localized and classified as MCF7. (**E**) Additional examples of FNs.

We further investigated the source of FNs. As shown in Table 2, the MSER localization module is unable to localize 71 MCF-7 cells, whereas the identification module misclassifies 22 MCF-7 cells. Therefore, the localization errors are responsible for more than 75% of the FNs. Majority of FNs encountered during identification are MCF-7 cells whose contrast, shape, and size differ from an average MCF-7 cell (See examples in Fig.4E). These two observations indicate that the convolutional layers of the CNN classifier were able to grasp the common features of MCF-7 cells for identification. As observed, the decoupled nature of this approach enabled us to trace back the missed detections of the CNN classifier and study them to gain insights into the discriminatory power of features that convolutional layers extract in classifying MCF-7 cells versus WBCs.

### B. Faster R-CNN Cell Detection

The decoupled cell detection approach performs the MCF-7 localization and identification in two steps. However, the sensitivity of this approach is being affected by the large number of FNs due to localization errors. We therefore explored a Faster R-CNN model, which integrates these two steps without having separate access to the output of the localization step or the input of the classification step. The Faster R-CNN model takes a brightfield image as input and outputs a bounding box (BB) for every MCF-7 cell present in the input image (Fig. 5). Two major modules underlie the Faster R-CNN model: (1) a localization module, *i.e.*, a Region Proposal Network (RPN) that discovers regions in the input image that are likely to include an MCF-7 cell, (2) a classification module, *i.e.*, Fast RCNN that labels the region proposals which essentially include an MCF-7 cell and estimates a BB that properly confines the cell.

**Figure 5.**
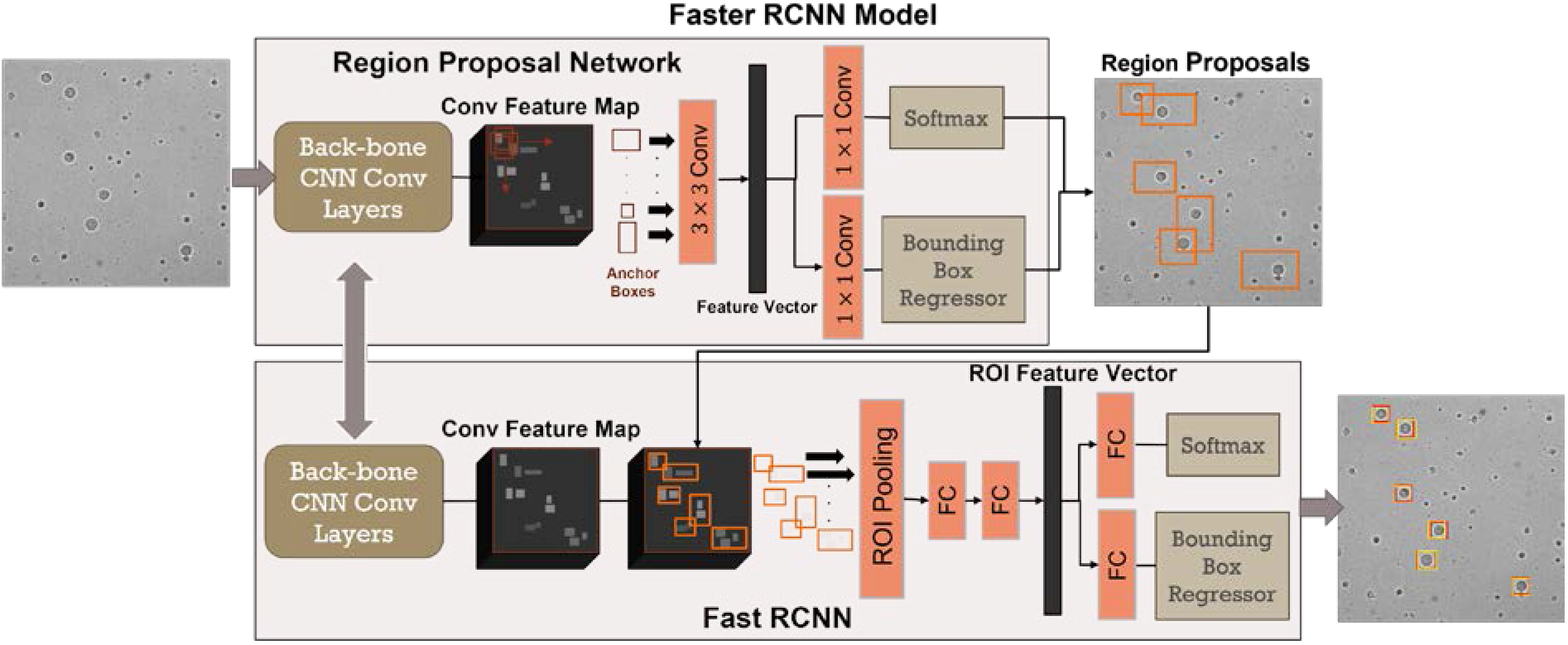
The Faster R-CNN cell detection approach. The Faster R-CNN detection model consists of two main integrated modules: (i) a Region Proposal Network that generates the region proposals (indicated by the orange bounding boxes in the image) that are most likely to include an MCF-7 cell, and (2) a Fast RCNN that labels the generated region proposals and outputs a bounding box for each detected MCF-7 cell. In the output image, the yellow boxes give the estimated bounding boxes assigned to the MCF-7 cell and the red boxes depict the corresponding ground truth.

#### Automated data set generation for training

For training a Faster R-CNN model, every image from the training dataset must be accompanied by a BB associated with each MCF-7 cell present in the image. The manual generation of BBs can be a tedious, subjective, and labor-intensive task. Here, we propose a technique for generating the BBs automatically; see Fig. 6. In this technique, the MCF-7 cell signatures in the FITC mask are first localized utilizing the circular Hough transform algorithm [45]. The FITC mask background is less noisy and complex than its corresponding brightfield image and, therefore, a better option to use for localization. For every localization, if not touching the image boundaries, the corresponding region in the DAPI mask is screened. If a signature exists in the screened region, then the corresponding localization is added to the MCF-7 binary mask. Lastly, for every blob in the MCF-7 binary mask, a 36 × 36 BB is generated and centered on that blob.

**Figure 6.**
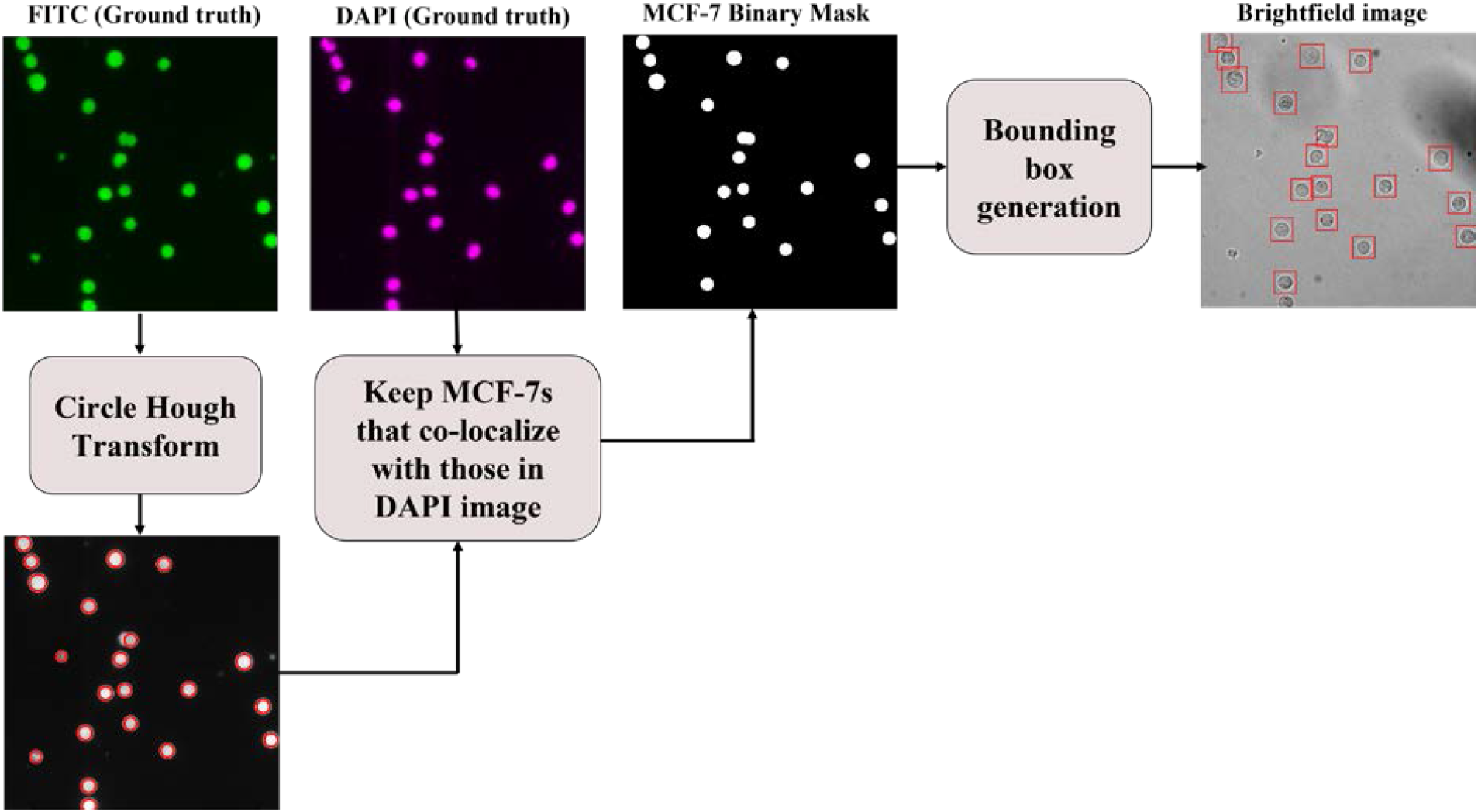
Dataset generation procedure for training the Faster R-CNN cell detection approach. For every brightfield image, the MCF-7 cells are first localized in the corresponding FITC mask using the circular Hough transform. Then, each localization, if not touching the image boundaries, is screened in the DAPI mask. If the localized region has a signature in the DAPI mask, then it is added to the MCF-7 binary mask. Finally, a bounding box is generated for every blob in the MCF-7 binary mask.

With this technique, 613 mixed population and pure cell images were annotated in less than 117 seconds, which is a significant improvement compared to manual annotation taking more than 100 seconds per image on average using MATLAB’s Image Labeler tool. Additionally, this technique produced equal-sized BBs that are centered on the cells; an outcome that is difficult to achieve manually. Consequently, the proposed automatic dataset generation algorithm is a fast, efficient and consistent technique to generate BB annotations for MCF-7 brightfield images.

#### Performance validation of the Faster R-CNN approach

For the Faster R-CNN cell detection approach, we utilized the same training images used in the decoupled cell detection approach. A total of 2,760 MCF-7 BBs were generated for the 355 primary training images. We employed the ResNet50 [46] as the back-bone CNN for the Faster R-CNN model. The negative and positive overlap ranges were set to [0,0.5] and [0.6,1], respectively. The model was trained for 30 epochs on an Nvidia TITAN RTX in 285 minutes. The model’s performance on the training images indicated an average Intersection-Over-Union (IoU) of 80.3% and a prediction score of 99.6%. The trained model achieved 99.6% training sensitivity and 99.18% training precision.

For the Faster R-CNN detection approach, the TPs, FNs, and FPs were defined in the same way as the decoupled approach, however they were interpreted differently due to the dissimilarities in the mechanism of the two approaches. All MCF-7 cells assigned a BB having *IoU* > 50% with respect to the annotation BB were counted towards TPs (Fig. 7A). An MCF-7 cell missing an estimated BB was considered a FN (Fig. 7B). There were two kinds of FPs as shown in Fig. 7 (C, D): (1) extra BBs assigned to a single MCF-7 and (2) non-MCF7 objects assigned a BB.

**Figure 7.**
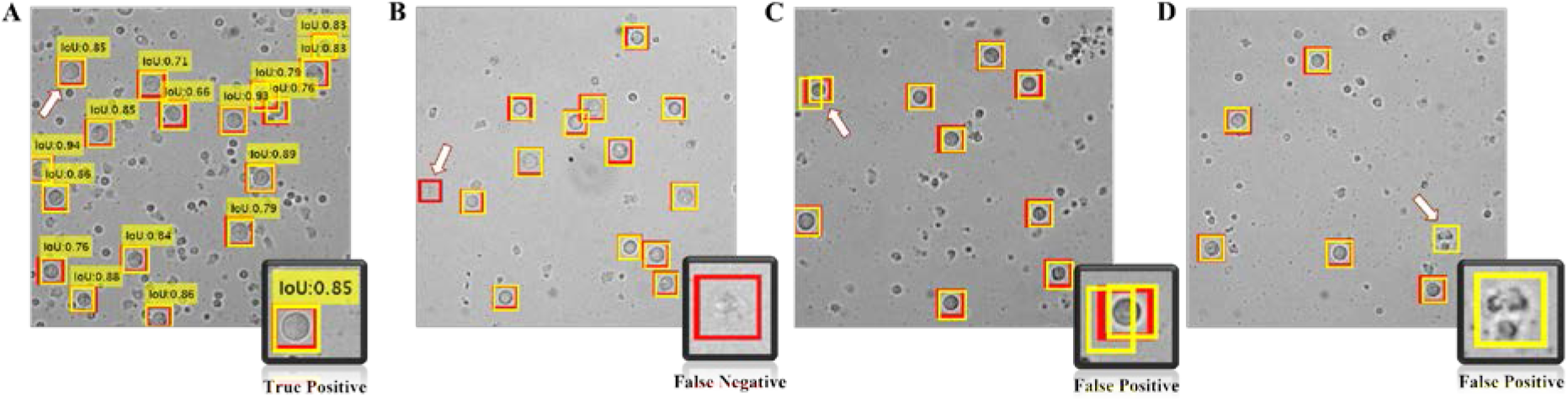
Representative examples of (A) true positive (B) false negative and (C, D) false positive from the Faster R-CNN approach. The arrow in each image and the bottom right inset highlights these examples. The yellow boxes give the estimated bounding boxes assigned to the MCF-7 cell and the red boxes depict the corresponding ground truth.

#### Performance comparison of the two approaches

As discussed in the previous sections, we developed two different frameworks that detect MCF-7 cells in the mixed cell population images. The first developed approach, *i.e*., decoupled cell detection, did not demonstrate acceptable sensitivity rate, which encouraged us to seek an alternative cell detection approach, *i.e.*, Faster R-CNN detection approach. Table 2 presents the statistical measurements and corresponding performance metrics of the two cell detection approaches. Both approaches exhibit a comparable precision, whereas the sensitivity of the Faster R-CNN approach is better than the decoupled approach, indicating that the Faster R-CNN approach is less likely to miss MCF-7 cell detections. Note that unlike the decoupled approach, we cannot trace back the missed detections (FNs) specifically to the localization or classification modules in the Faster R-CNN model.

Comparing the performance of both approaches indicate that the Faster R-CNN cell detection approach is preferred because it provides better overall performance in detecting MCF-7 cells in roughly half the time. Finally, the cell detection framework proposed by Wang *et al.* [36] is comparable to our decoupled cell detection approach in that it too is comprised of cell localization and cell identification modules. However, their work does not offer feature analysis, nor do they assess the performance of each module independently. Moreover, our proposed deployed detection model, i.e., Faster R-CNN model, outperforms Wang *et al.*’s detection framework.

### C. Impact of image intensity transformations on the Faster R-CNN approach

A major factor determining the effectiveness of a trained deep-learning model is how well it performs when exposed to images with variations in the background intensity and contrast, as well as intensity variations within cells and cell boundaries. Such variations are not uncommon in experimental images due to variability in cell sample preparation and imaging conditions (e.g., focus position, exposure times). To understand the effect of such variations on the performance of Faster R-CNN model, we performed image transformations and assessed the performance of the Faster R-CNN approach on the transformed images. In addition to sensitivity and precision, F1-score was used to evaluate the overall performance of the Faster R-CNN detection model in the presence of image transformations. F1-score is defined as the harmonic average of recall and precision, evaluating the overall changes in the performance of the detection model, when deployed on a dataset.

We performed image transformations and calculated the aggregate intensity distribution of a dataset by averaging the accumulated intensity distributions of all the images in the corresponding dataset. The intensity histogram of every image, *h*(*x*), was altered in the following three ways:

1. Shift the image intensity histogram to generate a brighter version of the image. The offset (*B*_1_) added to every pixel intensity is randomly drawn from a uniform distribution with bounds 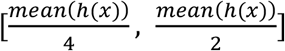. The altered image is modeled with ***I***_*Int*_(*x*, *y*) = ***I***(*x*, *y*) + *B*_1_ with corresponding *h*_*Int*_(***I***). This transformation increases the mean of *h*(***I***) by 25%-50%. The upper limit was set to avoid any pixel intensity saturation. The aggregate of all generated *h*_*Int*_(***I***) is represented by *H*_*I*_.
2. Increase the image contrast through stretching the image intensity distribution, *h*(***I***), around the mean (increase the intensity variance). For this purpose, the offset (*B*_2_) is chosen so that *h*(***I***) is centered in the [0, 255] range. This offset is added to every pixel intensity value. Afterwards, a random value (*A*_2_) is chosen from a uniform distribution with bounds [1.5, 2.5] and multiplied to every pixel value. This results in a darker version of the image with higher contrast. The altered image and its corresponding histogram are represented with ***I***_*IC*_(*x*, *y*) = *A*_2_ × (***I***(*x*, *y*) + *B*_2_) and *h*_*IC*_(***I***), respectively. By this transformation, the variance of *h*(***I***) rises by 125%-525%. The aggregate of all generated *h*_*IC*_(***I***) is represented by *H*_*IC*_.
3. Invert the pixel intensity values to generate the negative of the corresponding image. The altered image and its corresponding histogram are represented with ***I***_*N*_(*x*, *y*) = 255 − ***I***(*x*, *y*) and *h*_*N*_(***I***), respectively. The aggregate of all generated *h*_*N*_(***I***) is represented by *H*_*N*_.

In addition to the above three transformations that were performed on the original image data sets discussed in the previous sections, we also generated a new experimental image data set by altering the focus and exposure time during image acquisition. The aggregate intensity distribution for this new data set is denoted by *H*_*newexpt*_.

In Fig. 8A, we show the aggregate intensity distribution *H*_*TR*_ and *H*_*TS*_ for our original image data set that was used in training and testing respectively of the Faster R-CNN model. As expected, the intensity distributions have similar mean and variance indicating that the detection framework is trained on a similar distribution as the testing dataset. In contrary, as demonstrated in Fig. 8 (B, C, E), *H*_*I*_, *H*_*IC*_ and *H*_*newexpt*_ have different mean values compared to *H*_*TR*_. Additionally, Fig. 8C shows notable differences in the variance of *H*_*IC*_ and *H*_*TR*_. Note that Fig. 8D displays a mirroring effect between *H*_*TR*_ and *H*_*N*_. This property suggests that, in contrast to the original training images, the cells in the negative images appear brighter than the background.

**Figure 8.**
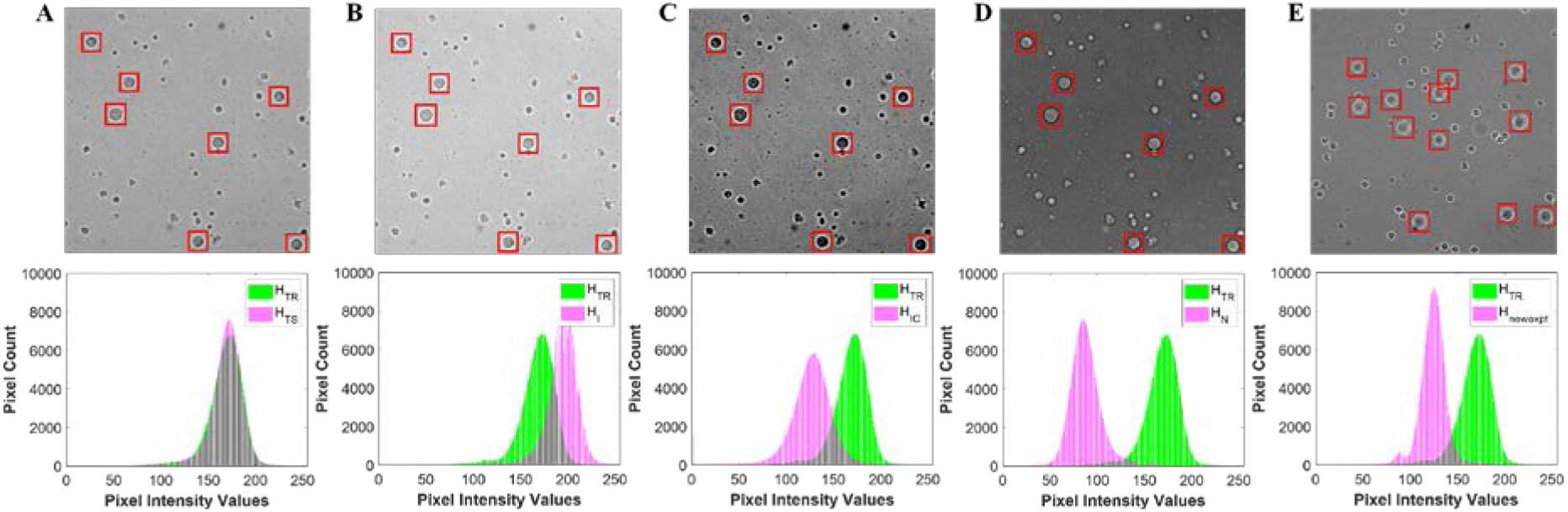
Impact of image transformations on the aggregate intensity distributions. (**A**) An original testing image and the corresponding aggregate intensity distribution, *H*_*TS*_ (**B**) Intensity altered image and the corresponding aggregate intensity distribution *H*_*I*_ (**C**) Intensity and contrast altered image and the corresponding aggregate intensity distribution *H*_*IC*_ (**D**) Negative image and the corresponding aggregate intensity distribution *H*_*N*_ (**E**) New experiment image and the corresponding intensity distribution *H*_*newexpt*_. Each of the altered intensity distributions are compared with the intensity distribution *H*_*TR*_ from the original training data set.

Table 3 compares the performance of the Faster R-CNN approach on the original data set, the altered versions of the original data set, and the new experimental dataset. As shown, the detection model manifests its best performance, *i.e*., achieves highest F1-score, on the original dataset. This observation is rather intuitively comprehensible since the original testing dataset has a similar distribution as the original training set, and so, the detection model is trained for the data in the original testing dataset. Taking the performance of the model on the testing dataset as the reference, for 25%-50% intensity variation and 125%-525% variance or equivalently contrast change, the F1-score was decreased to less than 0.1% and 0.2%, respectively. These observations indicate that the Faster R-CNN approach robustly responds to the intensity and contrast variations. Note that the degradation in the performance of the Faster R-CNN approach extends as the transformed intensity distribution further deviates from the intensity distribution of the original training set. Based on the preceding observations, it can be fairly concluded that the detection model extracts features that are relatively robust to intensity and contrast variations.

**Table 3:** The performance of the Faster-RCNN approach on the original testing and transformed datasets

| Dataset | Histogram | Sensitivity | Precision | F1-score |
| --- | --- | --- | --- | --- |
| *Original testing image set* | $H_{TS}$ | 99.1% | 99.8% | 99.45% |
| *Intensity-altered image set* | $H_I$ | 98.9% | 99.9% | 99.40% |
| *Intensity- and contrast-altered image set* | $H_{IC}$ | 99.0% | 99.5% | 99.25% |
| *Negative image set* | $H_N$ | 98.8% | 71.1% | 82.70% |
| *New experimental image set* | $H_{newexpt}$ | 99.4% | 97.3% | 98.34% |

For the negative dataset, the precision noticeably decreased (−28.7%), causing the F1-score to drop 16.8% which shows that the model is considerably less robust to transformations that reverse the intensity map of the objects with respect to the background. However, when we retrained the Faster R-CNN model on training images and their negative versions and test the retrained detection model on the negative of primary testing images, the performance recovered to 99.1% sensitivity, 97.3% precision and F1-score of 98.2%. Thus, the detection model tends to show vulnerability towards contrast reversal transformations. However, if trained on both the image and its negative version, the detection model manages to learn features of the cell that are independent of the cell versus background contrast.

### D. Application of deep learning techniques to live patient-derived CTC detection

Our results show that the DL-based Faster R-CNN model can successfully classify live breast cancer cells in a background of white blood cells. This outcome parallels the success that deep learning models are achieving in the fields of biomedical image analysis and clinical diagnostics [47, 48]. These advances have led to successful efforts in applying deep learning to detect patient-derived CTCs in fluorescently stained blood samples [34, 36]. Applying similar techniques for detecting live CTCs in unstained patient blood samples would open new opportunities (e.g., ex vivo expansion of CTCs for drug screening and molecular analysis). However, application of deep learning techniques for label-free detection of patient-derived CTCs presents challenges as we discuss below.

For supervised machine learning-based models, generation of valid ground truth data is essential. In the case of patient-derived CTCs, generation of ground truth images requires live-cell fluorescent markers that are specific to CTCs but do not stain blood cells. However, tagging CTCs with live-cell fluorescent labels has been a challenge in the field, although some progress is being made. For example, Wang *et al*. used Carbonic Anhydrase 1X PE-conjugated antibody along with a live cell dye, Calcein AM to detect live CTCs in metastatic renal cell carcinoma patients [36]. In another study, a group of near-infrared (IR) heptamethine carbocyanine dyes were used to identify viable CTCs recovered from prostate cancer patients [49]. These dyes were shown to actively tag cancer cells in xenograft models and have since begun to be used as live markers to further improve cancer prognosis and treatment efficacy [50]. Recently, positive selection approaches involving the use of cell surface markers such as HER2, EpCAM and EGFR have also been implemented to recover viable CTCs [51].

Another challenge for generating ground truth images is the heterogeneity of patient-derived CTCs. During the metastatic cascade, primary tumor cells which are of epithelial origin become migratory acquiring a mesenchymal phenotype [52–54]. Studies with cancer patient blood show that some of the isolated CTCs have high epithelial marker expression while others have high mesenchymal marker expression or a combination of both [55–58]. This heterogeneity in marker expression requires multiple fluorescent labels to tag live CTCs and capture the diversity present in patient blood and generate ground truth data.

Finally, machine learning-based approaches often require large data sets for training. Since CTCs are present in low counts in patient blood, generating large, annotated data sets requires access to many patients’ blood samples. Collecting blood samples from a large patient population can be a time-consuming, expensive, and a logistically challenging task.

A potential avenue to address these challenges in label-free live detection of CTCs is to develop robust DL-based approaches to detect blood cells in patient samples and score the remaining non-blood cells as prospective CTCs. This negative selection-based approach will at least help in quickly screening which patient blood samples are of most interest for further scrutiny.

## 5. Conclusions

In this work, we proposed an automated framework for label-free detection of MCF-7 breast cancer cells in a background of WBCs in brightfield images. An effective Faster R-CNN-based detection model was developed for detecting MCF-7 cells in the acquired brightfield images. The proposed model demonstrated 99.1% sensitivity and 99.8% specificity, and an average IoU of greater than 80%. The MCF-7 cell detection model analyzed each brightfield image in less than 0.3 seconds, more than 300 times faster than a human labeler. Also, we introduced a novel fully-automated technique for training set generation.

Additionally, we conducted multiple studies to investigate the discriminatory features that an effective CNN derives and employs to differentiate MCF-7 cells and WBCs. These studies showed that the size of the cells was the main distinctive feature that the CNN used to distinguish MCF-7 cells from WBCs. However, in the absence of the size feature, the CNN was still capable of learning other features to perform the identification task, with an acceptable, yet decreased, accuracy level.

Finally, we examined the performance of the detection model in the presence of numerous image intensity transformations. The results demonstrated that for intensity and contrast variations, the F1-score of the detection model was reduced by less than 0.2%. Therefore, the MCF-7 cell detection model was sufficiently robust with respect to intensity and contrast image transformations. These observations inform that the detection model uses intensity- and contrast-invariant features to perform the detection task.

The results from this study indicate that deep learning approaches could be potentially applied to detect CTCs in the blood of cancer patients. In the future, challenges related to live-cell and multi-marker fluorescent labeling of patient-derived CTCs and community-wide access to large, labelled datasets need to be addressed.

## Acknowledgements

This work was funded by the Cancer Prevention and Research Institute of Texas (Grant no. RP190658).

